# Iterative transfer learning with neural network for clustering and cell type classification in single-cell RNA-seq analysis

**DOI:** 10.1101/2020.02.02.931139

**Authors:** Jian Hu, Xiangjie Li, Gang Hu, Yafei Lyu, Katalin Susztak, Mingyao Li

**Affiliations:** Department of Biostatistics, Epidemiology and Informatics, University of Pennsylvania Perelman School of Medicine, Philadelphia, PA 19104, USA; State Key Laboratory of Cardiovascular Disease, Fuwai Hospital, National Center for Cardiovascular Diseases, Chinese Academy of Medical Sciences and Peking Union Medical College, Beijing 100037, China; School of Statistics and Data Science, Key Laboratory for Medical Data Analysis and Statistical Research of Tianjin, Nankai University, Tianjin 300071, China; Departments of Medicine and Genetics, University of Pennsylvania Perelman School of Medicine, Philadelphia, PA 19104, USA

**Keywords:** single-cell RNA-seq, clustering, cell type classification, transfer learning, neural network

## Abstract

An important step in single-cell RNA-seq (scRNA-seq) analysis is to cluster cells into different populations or types. Here we describe ItClust, an **I**terative **T**ransfer learning algorithm with neural network for scRNA-seq **Clust**ering. ItClust learns cell type knowledge from well-annotated source data, but also leverages information in the target data to make it less dependent on the source data quality. Through extensive evaluations using datasets from different species and tissues generated with diverse scRNA-seq protocols, we show that ItClust significantly improves clustering and cell type classification accuracy compared to popular unsupervised clustering and supervised cell type classification algorithms.

## Background

Only recently has the single-cell RNA sequencing (scRNA-seq) technology matured. Emerging scRNA-seq studies have transformed our understanding of cell biology and human disease. An important step in scRNA-seq analysis is to identify cell populations or types by clustering [1]. Knowledge in cell types can reveal cellular heterogeneity across tissues, developmental stages and organisms, and improve our understanding of cellular and gene function in health and disease. Despite the unprecedented power of scRNA-seq, the high-dimensionality and inherited high level of technical noise are major hurdles for cell type identification. Popular scRNA-seq clustering methods such as Louvain’s method [2], SIMLR [3], and SC3 [4] may perform poorly for data with closely related cell types or low sequencing depths. Although denoising methods such as SAVER [5] and DCA [6] can provide more accurate gene expression estimates and help clustering, these methods are unsupervised, and cannot utilize cell type-specific gene expression information.

Since a large amount of well-annotated scRNA-seq datasets are already available, many state-of-the-art methods start to utilize information in these well-annotated datasets to aid cell type identification in new data. For example, scmap [7] projects cells in a target dataset to a space determined by highly informative genes selected from a well-labeled source dataset, and then assigns cell identities for cells in the target data based on their correlation with average cell type-specific gene expression in the source data. Moana [8] trains a support-vector-machine (SVM) with a linear kernel on principal component analysis (PCA)-transformed labeled source data, which are subsequently used to cluster cells in the target data. Seurat 3.0 [9] classifies cells in target data by finding anchor cell pairs between a well-labeled source dataset and the unlabeled target dataset. Both scmap and Moana learn cell type-specific gene expression information only in the source data, but ignore useful information in the target data, thus are vulnerable to batch effect between the source and target data. Although Seurat 3.0 utilizes information both in the source and target data through the identification of anchor pairs, it does not specifically utilize cell type label information in the source data.

An ideal approach for cell type identification should be able to utilize cell type-specific gene expression information both in the well-labeled source data and the unlabeled target data. Since the source and target data provide different amount of cell type-specific gene expression information, it is desirable to use a data-driven approach to determine the contribution of each data type in analysis. Transfer learning, a machine learning method that focuses on storing knowledge gained while solving one problem and applying it to a different but related problem, suits perfectly for this purpose. Borrowing this idea, we developed ItClust, a supervised machine learning method that takes advantage of cell type-specific gene expression information learned from source data, to help clustering and cell type classification on newly generated target data. Unlike the unsupervised Louvain’s method [2], which requires users to specify ‘resolution’ to determine the number of clusters, ItClust is able to automatically determine the number of clusters in the target dataset. It is also superior to existing supervised classification methods in that cell types that are missing in the source data can be well separated in the target data.

## Results

### Overview of the method and evaluation

ItClust requires two input datasets, a source dataset that includes cells with well-annotated cell type labels, and a target dataset that includes cells that need to be clustered and annotated. **Fig. 1** shows an overview of the ItClust algorithm. ItClust starts from building a source network to extract cell type-specific gene expression signatures from the source data. This step enables initializing the second, which is the target network, with parameters estimated from the source network. The initialized target network is then further trained using cells in the target data to fine-tune the parameters, so that cell type-specific gene expression signatures in the target data are captured. This step is critical when there is strong batch effect between the source and target data, or when the source data have poor quality. Once fine-tuning is finished, the target network then returns clustered cells in the target data. As a data-driven approach, ItClust utilizes information from both the source and target data for clustering. This is different from prediction based methods such as Moana [8], which only uses information in the source data to build the prediction model.

**Figure 1.**
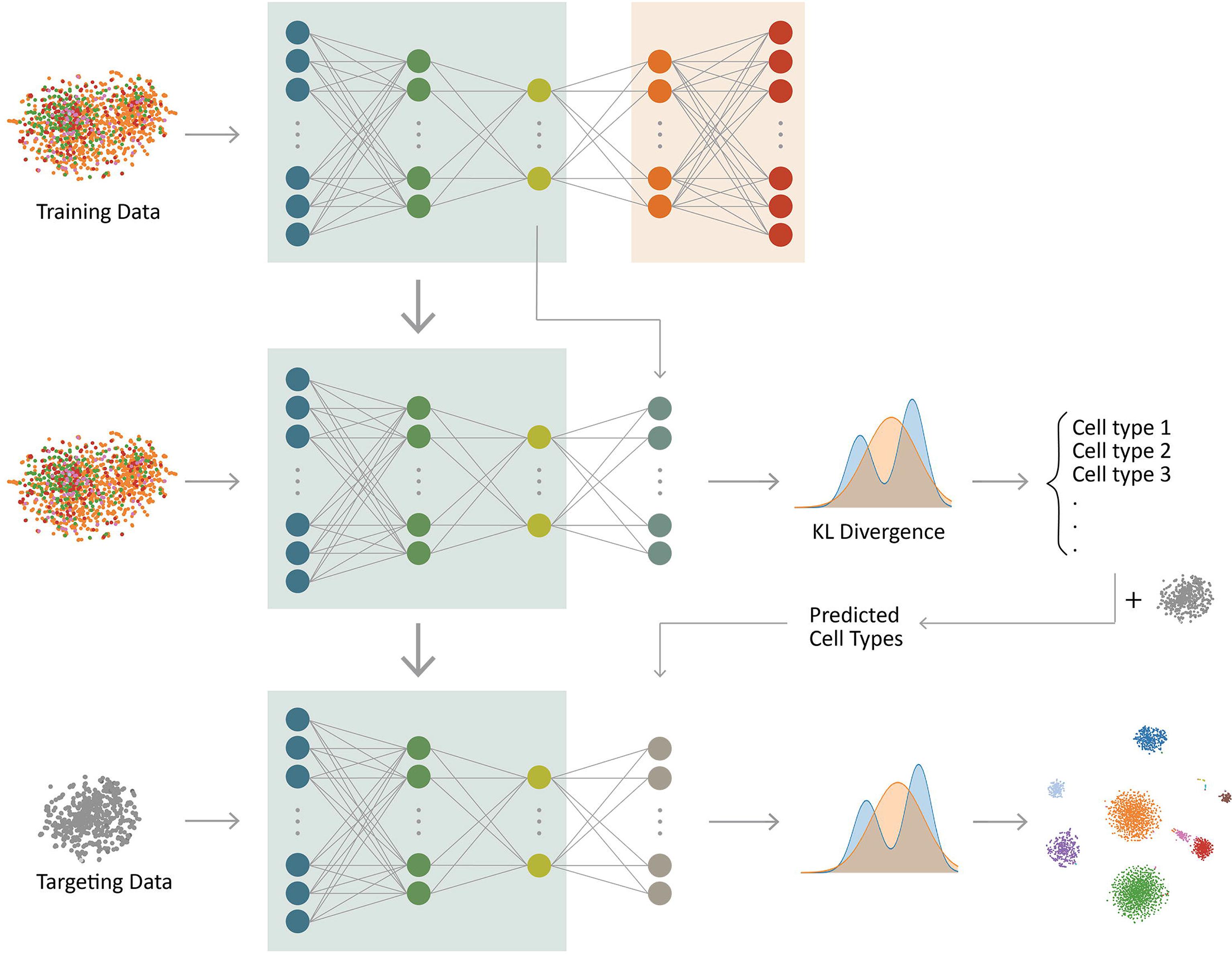
Overview of the ItClust framework. ItClust involves four steps: (1) parameter initialization for the source network with a stacked autoencoder; (2) parameter optimization (i.e. clustering) for the source network; (3) parameter initialization for the target network with information transferred from the source network, and (4) parameter optimization for the target network.

To showcase the strength and scalability of ItClust, we analyzed multiple scRNA-seq datasets from different species and tissues generated with different scRNA-seq protocols (**Supplementary Table 1**).

The performance of ItClust was compared with two unsupervised clustering methods including Louvain [2] and DESC [10], and three supervised cell type classification methods including Seurat 3.0 [9], Moana [8], and scmap [7]. We also compared ItClust with SAVER-X [11], a transfer learning algorithm for gene expression denoising. Our results show that ItClust consistently performs better than these existing methods in clustering and cell type classification.

### Comparison with unsupervised clustering methods

To show that incorporating cell type-specific gene expression information from well-labeled source data helps clustering in target data, we compared ItClust with Louvain and DESC, two unsupervised clustering algorithms. Louvain is a graph-based clustering method that has shown popularity in scRNA-seq analysis. DESC is an unsupervised neural network based clustering method that is effective in removing batch effect while maintaining high clustering accuracy. Since both are unsupervised, these two methods cannot utilize prior cell type information from a well-labeled source dataset. To make a fair comparison, we also compared ItClust with SAVER-X, a neural network based method originally designed for denoising of gene expression in scRNA-seq data. SAVER-X starts from extracting gene expression signatures from a source dataset and then denoises the unique molecular identifier (UMI) counts in target data with the learned prior knowledge on gene expression. After the data are denoised, clustering analysis can then be performed using algorithms such as Louvain. SAVER-X does not specifically utilize cell type label information in the source data.

We analyzed four publicly available datasets on human pancreatic islets generated using Fluidigm C1 ([12]; Lawlor data), SMART-seq2 ([13]; Segerstolpe data), CEL-seq ([14]; Grun data), and CEL-seq2 ([15]; Muraro data), respectively, as the target data. The human pancreatic islet data from Baron *et al*. [16], generated using InDrop, was used as the source data since this dataset contains a large number of cells from 14 well annotated cell types. For SAVER-X, the denoising performed using their pretrained model that includes both mouse and human pancreatic islet data. We first analyzed the four target datasets individually, and then combined them together to form a dataset to test whether ItClust is robust in the presence of batch effect in the target data. Cell type labels from the original publications were treated as the ground truth. Although the source and target data are both from the same species and the same tissue, they were generated through different scRNA-seq protocols, which result in strong batch effect between the source and target data and make transferring of cell type-specific gene expression knowledge difficult.

Since the performance of both the Louvain and DESC algorithms depends on ‘resolution’, a hyper-parameter that controls the number of clusters and needs to be specified by users, we tried a range of values from 0.2 to 2 with a step of 0.2. The Adjusted Rand Index (ARI) was used as the evaluation metric for clustering accuracy. **Fig. 2a** shows that across all four individual target datasets, the ARIs for Louvain, DESC, and SAVER-X vary substantially as the resolution parameter varies. In contrast, ItClust does not require the specification of the resolution parameter, and always had the highest or near the highest ARI even when comparing with the best performing resolution used by Louvain, DESC, or SAVER-X. For the combined dataset, the ARIs for Louvain, DESC, and SAVER-X dropped substantially because they tend to cluster cells from the same cell type but different datasets into different clusters, whereas ItClust maintained high clustering accuracy and is robust in the presence of batch effect in the target data (**Fig. 2b**). We note that unsupervised clustering algorithms such as DESC relies on batch indicator for standardization within each batch to remove batch effect, but supervised method such as ItClust does not need batch information in the target data.

**Figure 2.**
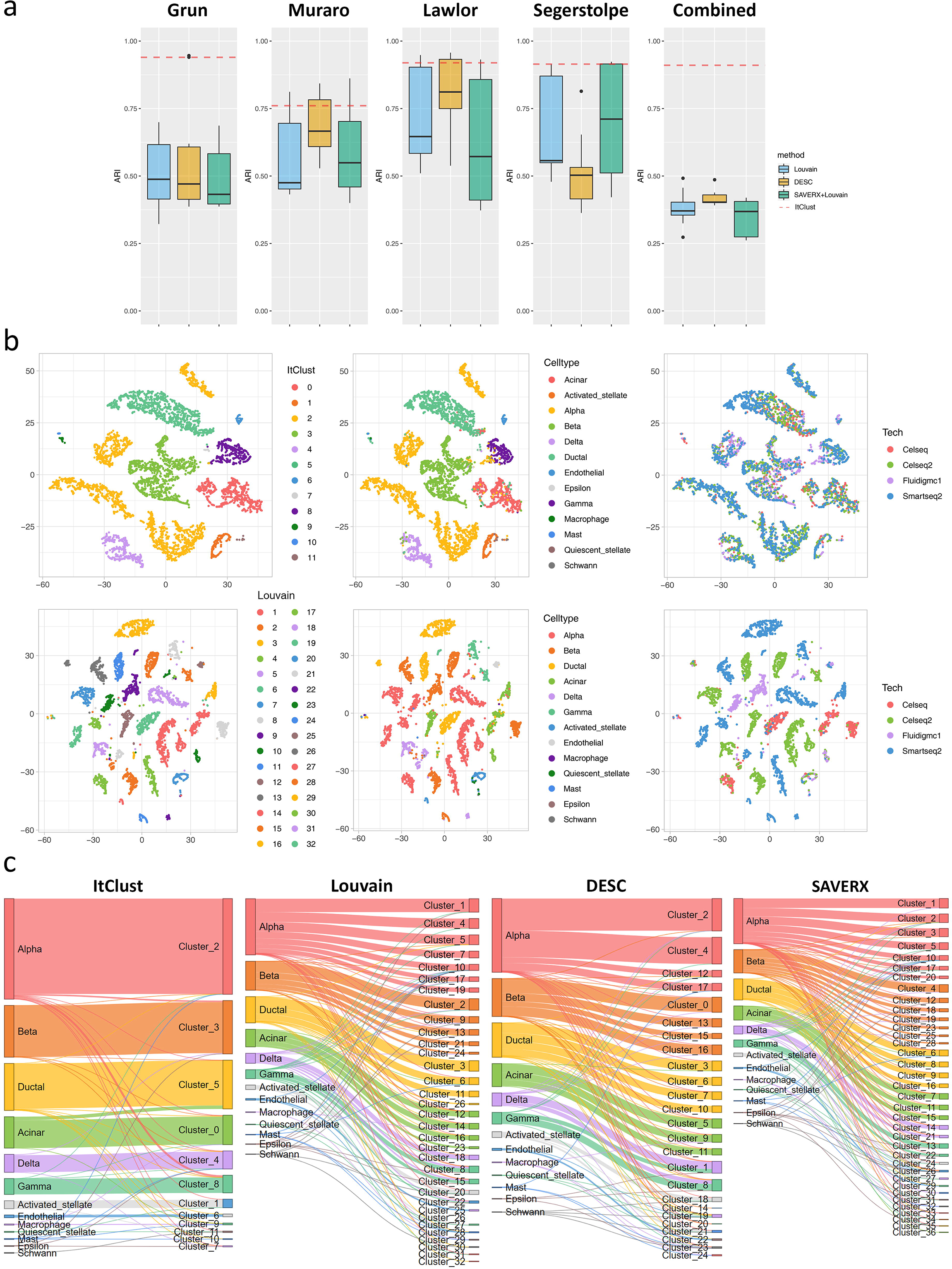
**(a)** The boxplots of clustering ARI of ItClust, Louvain [2], DESC [10], and SAVER-X [11] for the four human pancreatic islet datasets and the combined human pancreatic islet dataset [12–15]. Resolution of Louvain, DESC, and SAVER-X ranged from 0.2 to 2.0 with a step of 0.2. **(b)** The t-SNE plots for the combined human pancreatic islet dataset. The coordinates of the three plots in the first row are based on ItClust clustering result and colored by ItClust clusters, true cell types, and technical batches, respectively. The coordinates of the three plots in the second row are based on Louvain clustering result with resolution set to 2.0 and colored by Louvain clusters, true cell types, and technical batches, respectively. The minor cell types still mixed with other cells even when the resolution parameter was 2.0, suggesting the inability of separating minor cell types for Louvain. **(c)** The Sankey plots of ItClust, Louvain, DESC, and SAVER-X clustering results for the combined human pancreatic islet dataset. Resolution of Louvain, DESC, and SAVER-X was set to 2.0.

For Louvain, DESC, and SAVER-X, they can only separate major cell types even with increased resolution. As the resolution increases, Louvain and DESC tend to split major cell types into multiple sub clusters. However, the minor cell types still mixed with other cells even when the resolution parameter increased to a high value. In contrast, ItClust clearly revealed separate clusters for those minor cell types (**Fig. 2c**). The superior performance of ItClust over other methods is due to its use of cell type label information in the source data. Through training in the source data with well labeled cells, ItClust extracted gene expression signatures for each cell type, thus avoiding capturing information on batch effect, which makes it more sensitive to detect rare cells. Although SAVER-X also utilized prior information from the source data for denoising, cell type label information in the source data was not utilized, thus it did not show significant improvement in ARI after denoising compared to Louvain, which used the original gene expression data as the input.

### Comparison with supervised cell type classification methods

Next, we compared ItClust with supervised cell type classification methods. In addition to clustering, ItClust also provides a confidence score for each cluster, which indicates the degree of similarity of a cluster in the target data to an annotated cell type in the source data. Clusters with high confidence scores can be assigned with cell type names based on the corresponding annotations in the source data. For clusters with low confidence scores, they might represent cell types that are not present in the source data. In this case, we annotate their cell type names by known marker genes, which can be obtained from databases such as PangloDB [17] or CellMarker [18].

To evaluate ItClust’s performance for cell type classification, we assigned cell type names using approach described above, and compared ItClust with other supervised cell type classification methods based on accuracy, which is defined as the proportion of cells with correctly labeled cell type names. Among the pool of supervised classification methods, we selected Seurat 3.0 [9], Moana [8], and scmap [7] as our competitors. Seurat 3.0 is the most popular scRNA-seq analysis pipeline, and its cell type classification has shown good performance in recent publications. In Abdelaal *et al*. [19], SVM based methods are shown to have the best performance for intra-dataset classification. We chose Moana as a representative of the SVM family methods. Scmap is shown to be one of the best methods with a rejection option, which allows assigning cell types that are not present in the source data but are present in the target data as “unassigned”.

#### Within-species transfer

First, we considered the situation when source and target data are from the same species. We used the same four human pancreatic islet datasets analyzed previously as the target data, and the Baron human data as the source data. When considering each of the four target datasets separately, ItClust achieved the best performance in general, yielding the highest or near the highest classification accuracy (**Fig. 3a**). The next best performing method is scmap, followed by Seurat 3.0, and Moana.

**Figure 3.**
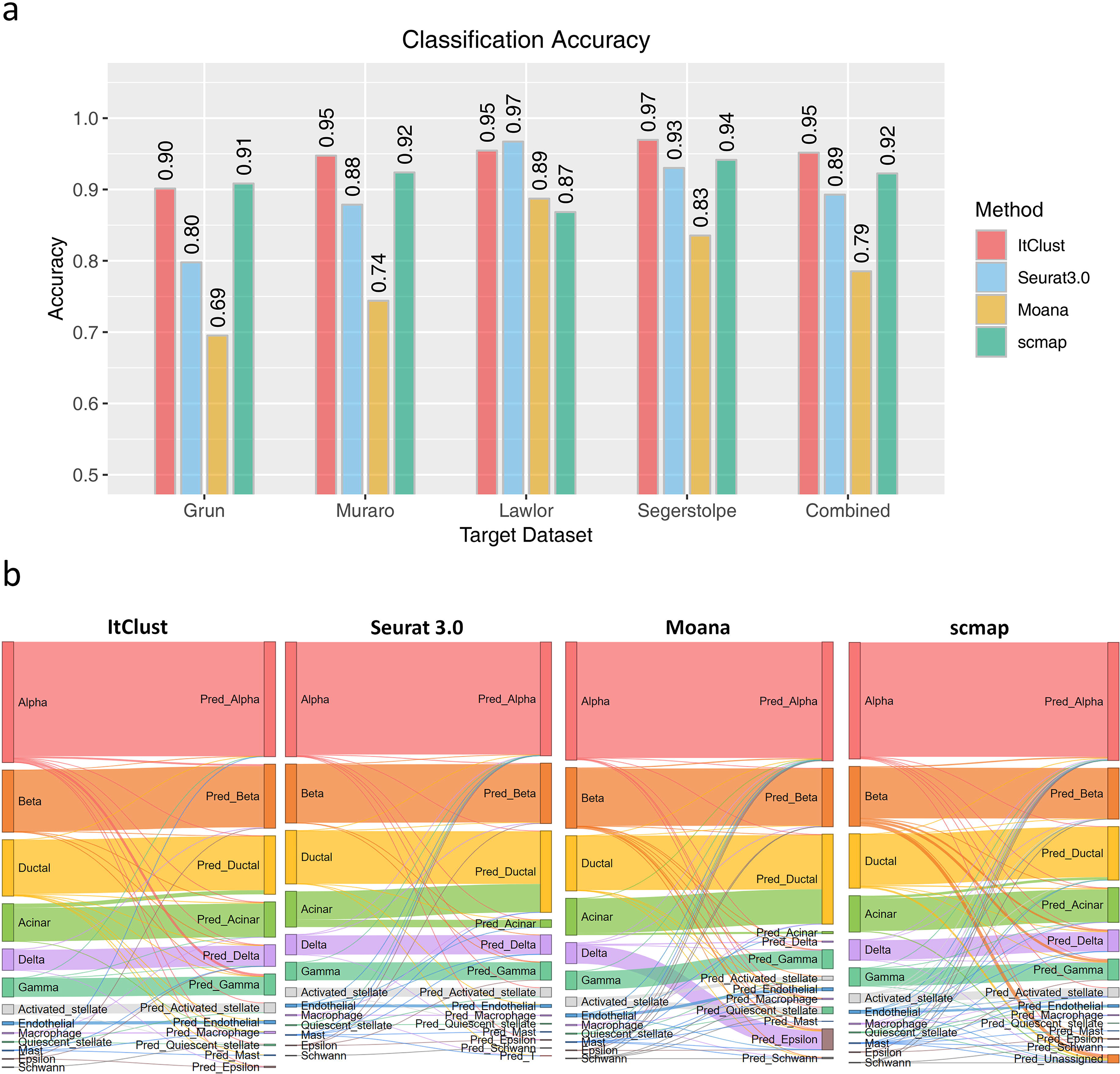
**(a)** The classification accuracies of ItClust, Seurat 3.0 [9], Moana [8], and scmap [7] for the four human pancreatic islet datasets and their combination [12–15], using the Baron human data as source [16]. **(b)** The Sankey plots of ItClust, Seurat 3.0. Moana, and scmap for the four human pancreatic islet datasets and their combination.

When the four target datasets were combined as a single target dataset, ItClust still achieved a high accuracy of 0.95 with each cluster corresponding to one cell type (**Fig. 3b**), which indicates its robustness to batch effect in the target data. Although Seurat 3.0 also achieved a high accuracy of 0.89, it tends to assign acinar cells to ductal cells (**Fig. 3b**). Seurat 3.0 uses an unsupervised strategy to pair anchor cells between two datasets, and cell type assignment in the target data is based on these anchors. By examining the overlap of mutual nearest neighbors for two cells in a pair, Seurat 3.0 also gives an anchor quality score that measures the similarity between the two cells. To further investigate why ItClust performed better than Seurat 3.0, we examined the anchors identified by Seurat 3.0. Among the 628 anchors for ductal cells in the source data, 189 anchors were mis-paired with an acinar cell in the target data, but these mis-paired anchors still have average anchor quality score of 0.85, which is significantly higher than the average anchor quality score for all 1,400 anchors (0.62). The mismatch of anchor cells is possibly due to the close relatedness of acinar and ductal cells in cellular differentiation because acinar, endocrine and ductal cells of the pancreas are all derived from a specific region of endoderm [20]. Since the anchor searching process is unsupervised, Seurat 3.0 would give a cell pair a high anchor score as long as they share similar gene expression pattern, although many of the genes that drive the similarity between the two cells are not marker genes for the corresponding cell type(s). ItClust avoids this issue by directly utilizing cell type information in the well-labeled source data to obtain the initial cluster centroids for the target data.

#### Cross-species transfer

Next, we considered a more challenging situation with the goal of transferring cell type knowledge learned from one species to a target dataset generated in a different species. First, we designed an experiment to transfer information from mouse kidney to human kidney. We used the Park *et al*. mouse kidney data [21] as the source data, and the human kidney data generated ourselves combined with another human kidney dataset generated by Young *et al*. [22] as the target data. Cells in the human data were clustered using DESC and annotated based on known marker genes [22] (**Supplementary Note** and **Supplementary Fig. 1**). As shown in **Fig. 4a**, ItClust achieved the highest cell type classification accuracy (0.87), which is much higher than the second best performing method Seurat 3.0 (0.69). Moana and scmap failed the task, yielding low accuracies of 0.20 and 0.19, respectively. Of note, Seurat 3.0 misclassified more than half of the macrophages (2,408 out of 3,566; 67.5%) as fibroblasts, whereas ItClust labeled 94.6% of the macrophages correctly **(Fig. 4b)**. To further verify these results, we selected marker genes for macrophage and fibroblast, and generated gene expression dot plots for the true cell types, and cell types predicted by ItClust and Seurat 3.0, respectively **(Fig. 4c)**. For the ItClust predicted macrophage cluster, known macrophage marker genes are expressed, whereas those marker genes for fibroblasts have low or no expression. In contrast, known macrophage marker genes have high expression in fibroblasts predicted by Seurat 3.0.

**Figure 4.**
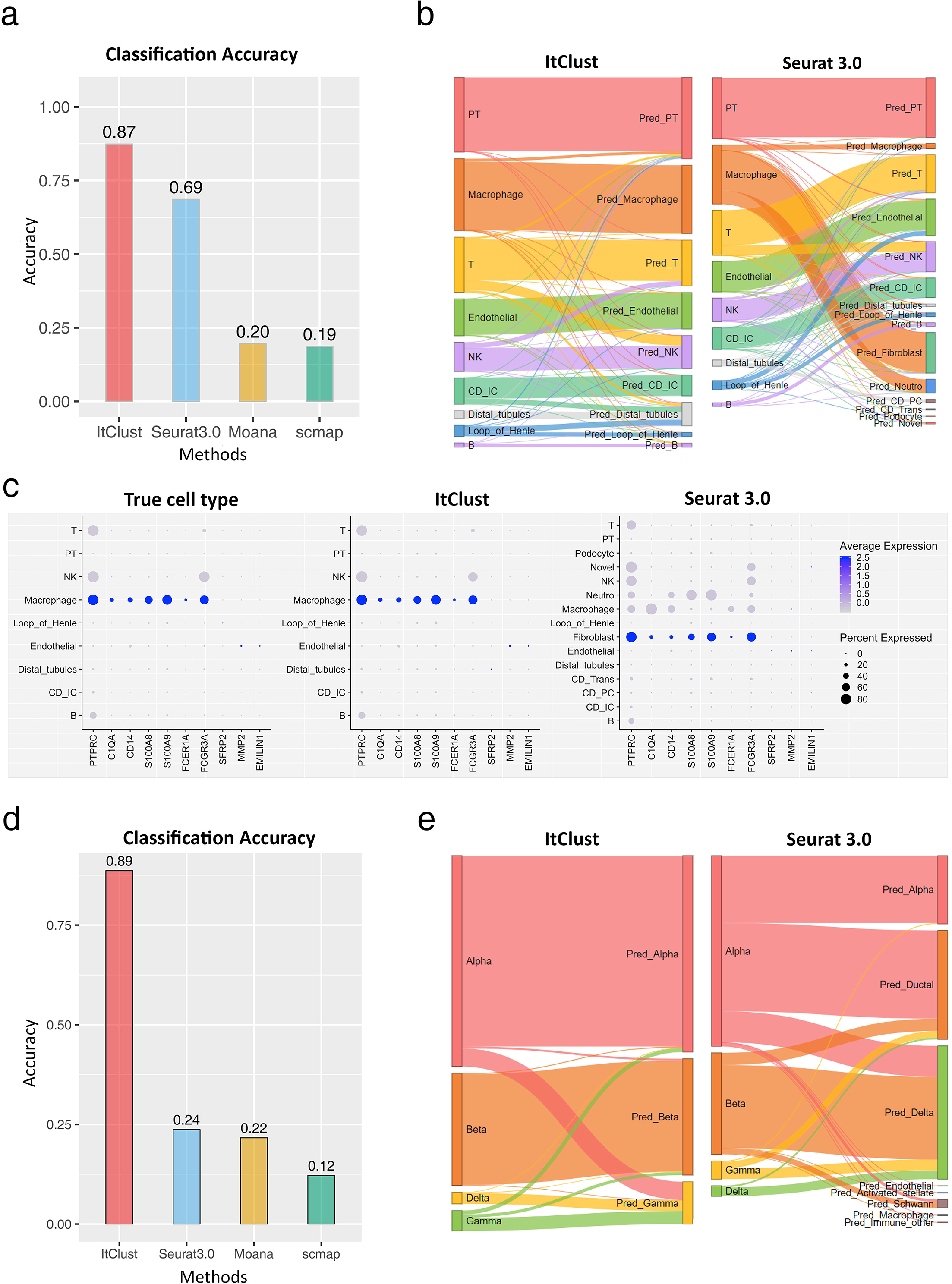
**(a)** The classification accuracies of ItClust, Seurat 3.0, Moana, and scmap for the human kidney dataset generated ourselves combined with another human kidney dataset generated by Young *et al*. [22], utilizing the mouse kidney data from Park *et al*. as source [21]. **(b)** The Sankey plots of ItClust and Seurat 3.0 for the combined human kidney dataset. **(c)** The dot plots of marker gene expression for macrophage and fibroblast for cells with true cell type labels, cells with cell type labels obtained from ItClust clustering, and cells with cell type labels obtained Seurat 3.0 clustering for the combined human kidney dataset. The seven macrophage marker genes (*PTPRC, C1QA, CD14, S100A8, S100A9, FCER1A* and *FCGR3A*) and the three fibroblast marker genes (*SFRP2, MMP2* and *EMILIN1*) were selected from Young *et al*. [22]. **(d)** The classification accuracies of ItClust, Seurat 3.0, Moana, and scmap for the human pancreatic islet dataset from Xin *et al*. [23], utilizing the mouse pancreatic islet data from Baron *et al*. as source [16]. **(e)** The Sankey plots of ItClust and Seurat 3.0 for the human pancreatic islet dataset from Xin *et al*. [23].

Encouraged by the above results, we further did a cross-species transfer analysis in pancreas. We used the Xin *et al*. [23] data generated from human pancreatic islets as the target data and the Baron mouse pancreatic islet data [16] as the source data. The target dataset only contains four cell types, which are all included in the source data. Despite the simplicity of the target data, the cross species transfer makes it difficult as there is a risk of transferring noise of gene expression signatures that are only present in mouse but absent in human. As shown in **Fig. 4d**, ItClust still achieved the highest classification accuracy (0.89), much higher than the second best performing method Seurat 3.0 (0.24). This is further confirmed by the Sankey plots shown in **Fig. 4e**. The significant advantage of ItClust over other methods is due to the iterative fine-tuning step, which enables the extraction of useful information that distinguishes cell types in the target data. The fine-tuning step is a critical advantage that differentiates ItClust from other supervised cell type classification methods.

#### Cell types that are not present in the source data

In many studies, the target data may contain cell types that are not present in the source data. To this end, we tested how unique cell types in the target data affect the performance of different methods. We made a human pancreas benchmark dataset based on the Baron human [16] and the Segerstolpe human data [13]. As shown previously, ItClust achieved the highest classification accuracy (0.97) in the Segerstolpe data with the Baron human data serving as the source. This provides an appropriate benchmark scenario that allows us to investigate how cell type classification accuracy varies as certain cell types are eliminated from the source data.

First, we considered a situation in which rare cell types are present in the target data but are missing in the source data. To do so, cells from epsilon, mast, macrophage, quiescent stellate, and schwann were eliminated from the Baron human data. As shown in **Fig. 5a**, ItClust still achieved a high accuracy of 0.96. The other methods also showed relatively high accuracies, with 0.92 for Seurat 3.0, 0.89 for Moana, and 0.94 for scmap. These results indicate that missing rare cells types in the source data had little impact for all evaluated methods.

**Figure 5.**
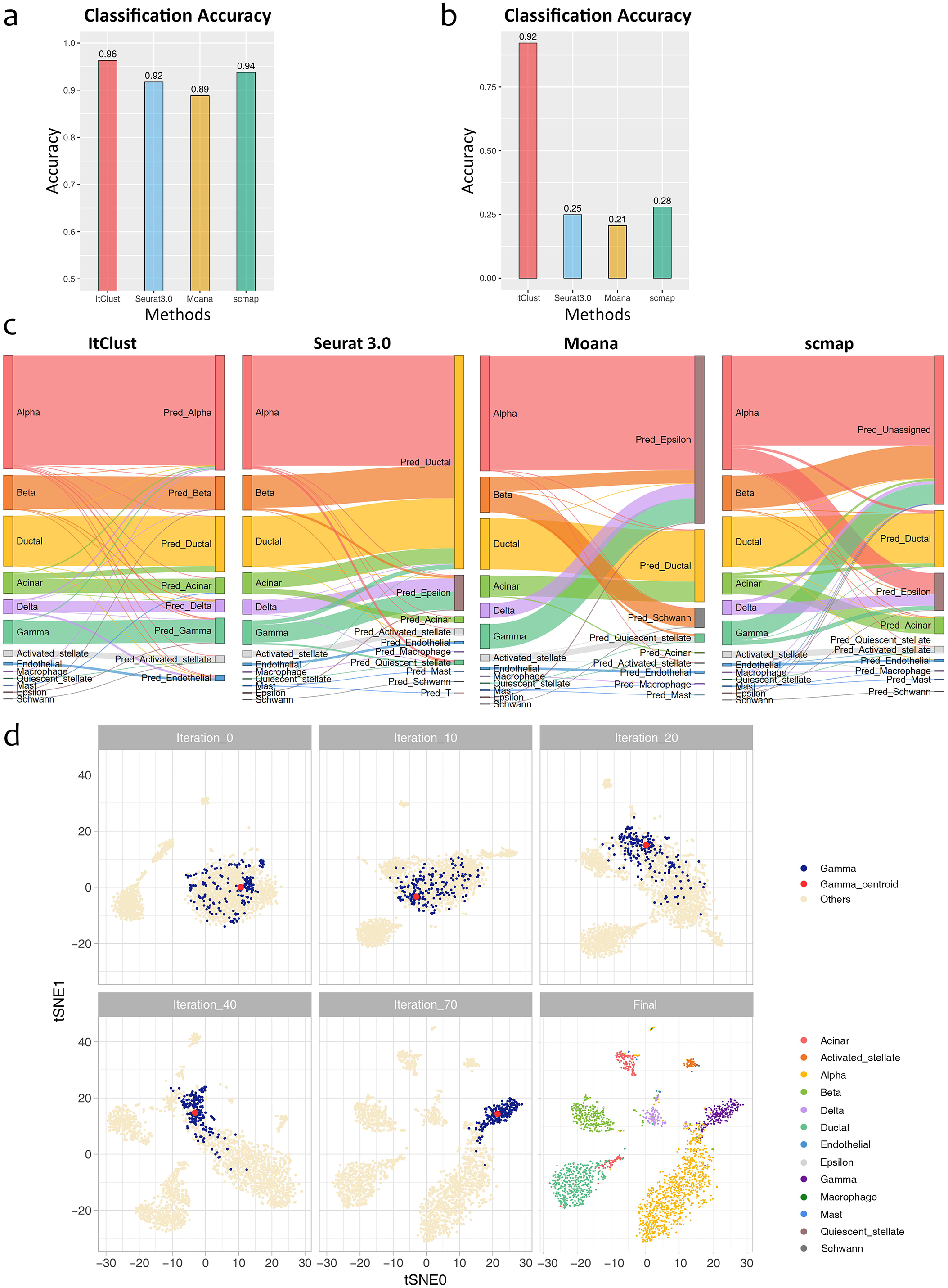
**(a)** The classification accuracies of ItClust, Seurat 3.0, Moana, and scmap for the Segerstolpe human pancreatic islet data, using the Baron human pancreatic islet data after eliminating minor cell types (epsilon, mast, macrophage, quiescent stellate, and schwann cells) and only keeping major cell types (alpha, beta, gamma, delta, ductal, acinar, activated stellate, endothelial, and T cells) as source in analysis. **(b)** The classification accuracies of ItClust, Seurat 3.0, Moana, and scmap for the Segerstolpe human pancreatic islet data, using the Baron human pancreatic islet data after eliminating major cell types (alpha, beta, gamma, and delta cells) and only keeping ductal, acinar, quiescent stellate, activated stellate, endothelial, macrophage, mast, epsilon, schwann, and T cells as source in analysis. **(c)** The Sankey plots of ItClust, Seurat 3.0, Moana, and scmap for the Segerstolpe human pancreatic islet data using the same reduced Baron human pancreatic islet data in (b) as the source. **(d)** The t-SNE plots of cells in the Segerstolpe human pancreatic islet data during the iterative fine-tuning in ItClust, using the same reduced Baron human pancreatic islet data in (b) as the source. The first five plots show the movement of gamma cells and the corresponding clustering centroid over iterations. The last plot shows the final clustering result for all cell types after the algorithm converges.

Next, we considered a more challenging situation by excluding four main cell types, including alpha, beta, gamma, and delta cells, from the source data. **Fig. 5b** shows that ItClust still achieved a high accuracy of 0.86 and was able to separate these four cell types correctly, although they were absent in the source data. In contrast, the accuracy of Seurat 3.0 dropped substantially to 0.25, since it assigned most of the four missing cell types as ductal (**Fig. 5c**). Scmap’s accuracy dropped to 0.28, and an additional 55.0% of the cells, including most of the alpha, beta, and gamma cells along with some acinar and delta cells, were classified as “unassigned,” and most of the delta cells (99 out of 127, 78.0%) were mis-assigned as epsilon cells. Moana, whose accuracy dropped to 0.21, also classified most of the unseen cells as eplison cells.

The reason that ItClust can separate these unseen cell types is that during the fine-tuning step, our model captured information of these unique cell types in the target data by updating the network parameters. Through iterative fine-tuning, these four cell types were gradually separated from the other cell types based on cell type-specific gene expression information gained from the target data. To better illustrate how the fine-tuning process works, we use gamma cells as an example. **Fig. 5d** shows a t-SNE plot of cells in the target data before fine-tuning. The blue dots represent those true gamma cells, and the red dot is the cluster centroid and the beige dots represent cells from the other cell types. Although we transferred cell type information from the source data to the model, because the gamma cells were absent in the source data, the initialized network was unable to separate those unseen gamma cells. Thus, in the t-SNE plot, gamma cells mixed together with the other unseen cell types initially. However, during the fine-tuning step, our model started to capture information of these unique cell types in the target data. Through iterations, the gamma cells were moving closer to the red centroid, which pulled the gamma cells away from cells that originate from cell types. After 70 iterations, the gamma cells were completely separated from the other cells. Similar patterns were also observed for alpha, beta and delta cells. These results indicate that for unseen cell types, if there is enough information in the target data, ItClust is able to utilize such information and cluster cells from those unseen cell types well.

#### Combined source data

In situations when it is hard to find one dataset that contains all needed cell types as the source, a simple solution is to combine multiple datasets to form a complementary dataset that includes as many cell types as possible, and then use this combined dataset as the source data. In this section, we demonstrate that ItClust is robust to batch effect in the combined source data and is able to extract useful information to help improve clustering and cell type classification. As shown in the previous section, when excluding the four main cell types (alpha, beta, gamma, and delta) from the Baron human source data, ItClust still had good classification accuracy on the Segerstolpe dataset. Here, we combined the reduced Baron human data with the Xin data, which only contain the four missing cell types, to generate a new combined source dataset. With this combined dataset as the source data, the overall classification accuracy of ItClust increased from 0.92 to 0.96 (**Fig. 6a**). The Sankey plot also shows that alpha, beta, gamma, and delta cells were all classified better (**Supplementary Fig. 2**). The classification accuracies for Seurat 3.0, Moana and scmap also improved when using the combined source data as the training data, but their accuracies are still lower than ItClust.

**Figure 6.**
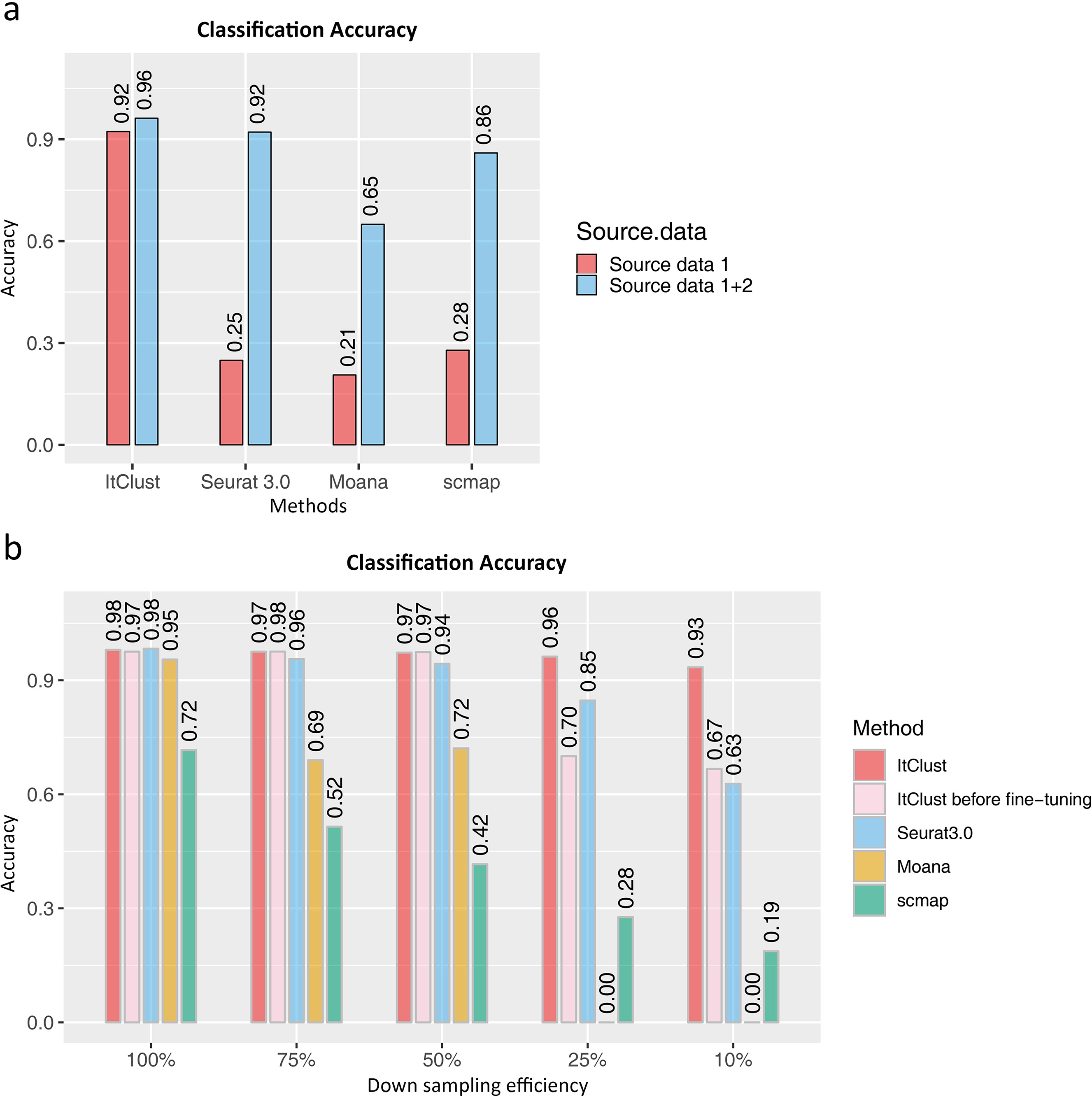
**(a)** The classification accuracies of ItClust, Seurat 3.0, Moana, and scmap for the Segerstolpe human pancreatic islet data, using different source datasets as input. Source data 1 is the reduced Baron human pancreatic islet data as in Figure 5(b) and source data 2 is the Xin human pancreatic islet data, which only include alpha, beta, gamma, and delta cells. **(b)** The classification accuracies of ItClust before and after fine-tuning, Seurat 3.0, Moana, and scmap for the macaque retina data across different down-sampling efficiencies. Cells from macaques 1, 2, and 3 were used as the source data, and cells from macaque 4 were used as the target data [24].

#### Read depth down sampling experiments

Many scRNA-seq studies suffer from low sequencing depth although they can easily generate a large number of cells. To evaluate the performance of ItClust when the source data have low sequencing depth, we performed read depth down sampling experiments using a scRNA-seq dataset on retina bipolar cells from four macaques [24]. We merged cells from the first three macaques and used the merged data as the source data, and cells from the fourth macaque as the target data. Since the source and target data are from the same study and were generated using the same protocol, ItClust, Seurat 3.0, scmap, and Moana all achieved high classification accuracy, thus providing an appropriate testing case for down sampling evaluation.

To do read depth down sampling on the source data, we set the sampling efficiency at 75%, 50%, 25%, and 10%, respectively [5], with lower efficiency corresponding to lower sequencing depth. As shown in **Fig. 6b**, as efficiency decreases, the accuracy of all methods declined. For example, when the sampling efficiency is 10%, the accuracies of both Seurat 3.0 and scmap dropped substantially, 0.98 to 0.63 for Seurat 3.0, and 0.72 to 0.19 for scmap, but ItClust only dropped slightly from 0.98 to 0.93. Moana failed to predict any cells when the efficiency dropped to 25% or lower because the quality of the source data is too low and Moana failed to build a prediction model. Compared to Seurat 3.0, ItClust is more robust to decline in read depth and less sensitive to the quality of the source data. This is because the anchor pairs identified by Seurat 3.0 depend on the quality of both the source and target data. When the source data quality is low, Seurat 3.0 has difficulty finding trustworthy anchor pairs. Since ItClust extracts cell type-specific gene expression in the target data through fine-tuning, it still gave high classification accuracy by utilizing more information in the target data. **Fig. 6b** shows that the fine-tuning step in ItClust significantly improved the classification accuracy when efficiency is as low as 25% and 10%. The ability to extract cell type-specific gene expression signatures from both the source and target data makes ItClust flexible.

#### Cell type assignment

In this section, we leverage cell type information in the source data to help cell type assignment in the target data. We define a confidence score that measures how confident that cells in a given cluster in the target data are similar to a cell type that is present in the source data (**Fig. 7a**). First, we consider the human pancreas data in the within species transfer section in which the source and target data include the same cell types. As shown in **Fig. 7b**, all clusters have high confidence scores, indicating high similarity between the source and target data. As another example, we calculated the confidence scores for the human pancreas data in which the four major cells were excluded from the source data. **Fig. 7c** shows that clusters 1 and 2 have relatively high confidence scores of 0.96 and 0.94, respectively. These two clusters correspond to activated stellate cells and ductal cells in the original source data. Therefore, we can confidently assign them as activated stellate cells and ductal cells for cells in the target data. For the other clusters with low confidence scores, we would encourage users to look at expression levels for known marker genes or use algorithms such as ACTIONet [25] for automatic cell type annotation.

**Figure 7.**
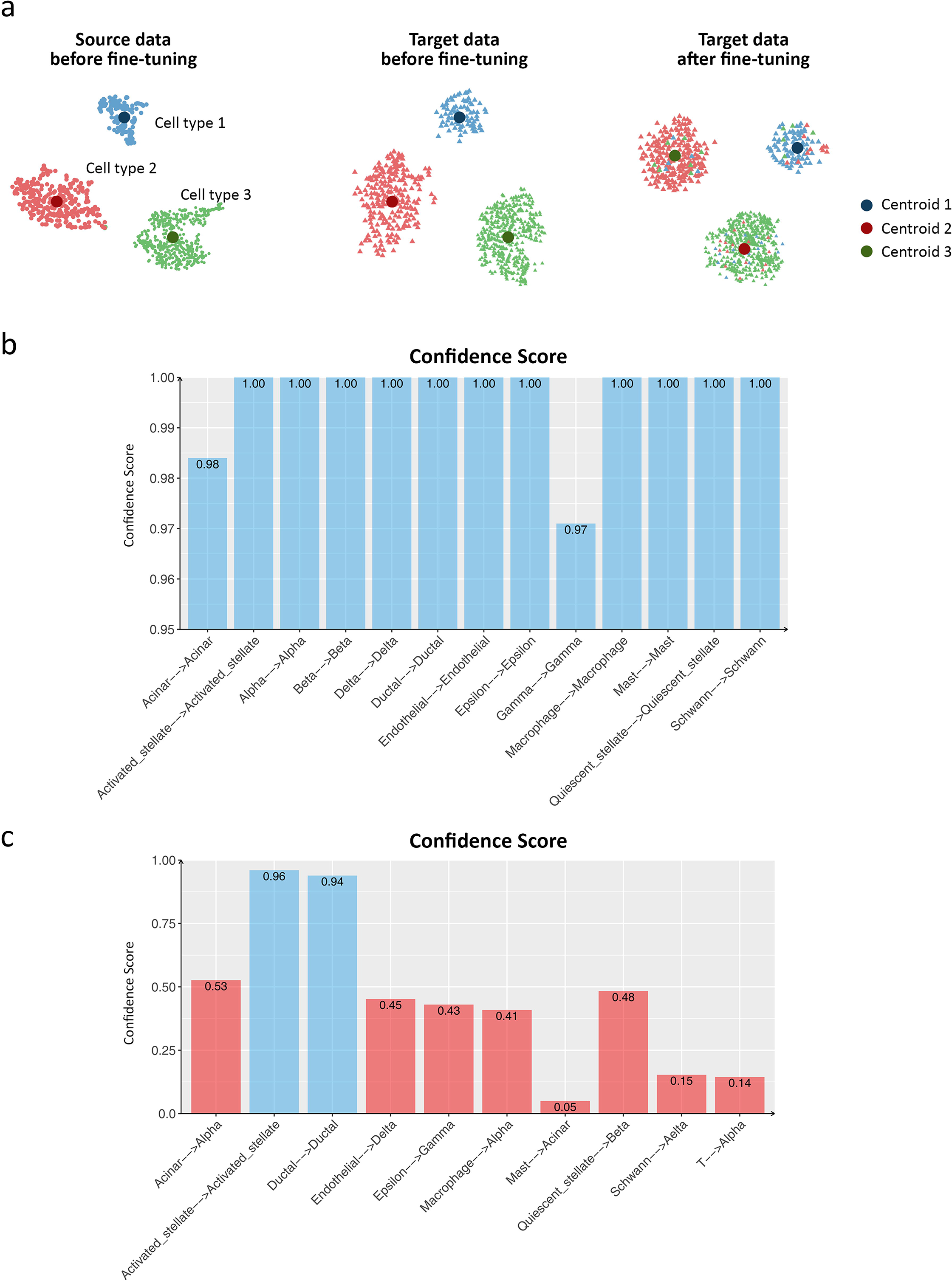
**(a)** Illustration of the confidence score calculation for cell type assignment in ItClust. Assume the source data only contain three distinct known cell types. Before fine-tuning on the target data, ItClust calculates and records the cluster centroid for each cell type in the source data. Then, ItClust uses these known cell type centroids as initial centroids to cluster cells in the target data. After fine-tuning is finished, the locations of the centroids are updated. Meanwhile, cells assigned to each centroid are updated. For example, most of the red cells in the target data are initially assigned to the centroid 2 that corresponds to cell type 2 in the source data, but switched to centroid 3 corresponding to cell type 3 in the source data after fine-tuning, as the location of these two centroids are changed over iterations. In contrast, the blue cells in the target data are consistently assigned to centroid 1 corresponding to cell type 1 in the source data before and after fine-tuning. Therefore, the blue cells in cluster 1 have a high confidence score to be assigned as cell type 1 and the red cells in cluster 3 have a low confidence score to be assigned as cell type 3. **(b)** The confidence score for each cluster for the Segerstolpe human pancreatic islet data, using the Baron human pancreatic islet data as the source. **(c)** The confidence score for each cluster for the Segerstolpe human pancreatic islet data, using the reduced Baron human pancreatic islet data when four major cell types (alpha, beta, gamma, and delta cells) were eliminated as the source data. For this benchmark dataset, we know the true cell type labels for both the source and target data, thus we can compare the cell types in the source and target data to examine if they are assigned to clusters corresponding to the same centroid. If the corresponding cell types for cells in the source and target data are the same, the confidence score should be close to 1. The x-axis labels in (b) and (c) indicate the change of cell type label assigned to the same centroid from the source data to the target data.

## Discussion

Clustering and cell type classification are important steps in scRNA-seq analysis. As more and more scRNA-seq datasets are becoming publicly available, supervised cell type classification methods that utilize external well-annotated data start to gain popularity over unsupervised clustering algorithms. However, a shortcoming of existing supervised methods is that their performance is highly dependent on quality of the source data, and the similarity between the source and target data. When strong batch effect between the source and target data exists or the quality of the source data is low, the performance of these methods drops drastically. Furthermore, most supervised classification methods assume cell types in the target data are similar or are a subset of cell types that are present in the source data. Although some methods, such as scmap, have a rejection option to allow unassigned cells in the target data, in many situations, the cells classified as “unassigned” are due to failure of correctly assigning them to existing cell types in the source data.

To address these limitations, we developed ItClust, a supervised clustering algorithm that employs a transfer learning framework. ItClust borrows idea from existing supervised cell type classification algorithms, but also leverages information in the target data to make it less dependent on the source data quality. This flexibility allows ItClust to utilize cell type-specific gene expression information from well-annotated datasets to help clustering without the need to specify resolution, a hyper-parameter that is required by unsupervised clustering algorithms such as Louvain and DESC. After clustering, ItClust also provides a confidence score for each cluster to assist cell type assignment. Compared to supervised classification methods, ItClust can utilize information extracted from both source and target data, making it robust when the quality of the source data is low. ItClust is also able to separate unseen cell types into different clusters, thus facilitating downstream analysis of those unseen cell types.

ItClust has been extensively tested using datasets from different species (mouse, macaque, and human), different tissues (pancreas, kidney, and retina) generated using nine different scRNA-seq protocols (10X, inDrop, DropSeq, SeqWell, CelSeq, CelSeq2, SmartSeq2, Fluidigm C1, and Smarter). Comparison of ItClust with other unsupervised clustering methods showcased that it always achieves high ARIs without the need of tuning any hyper-parameters such as resolution. The comparison with popular supervised cell type classification methods, i.e., Seurat 3.0, Moana, and scmap, also supported ItClust’s consistent high performance in all evaluated scenarios.

The success of ItClust comes from its unique way of information extraction and integration. Different from other methods, ItClust first extracts information from the source data by considering genes that are highly variable in the target data. This step ensures that all the transferred expression patterns are useful for separating cell types in the target data. These expression patterns are further updated based on information in the target data through iterative fine-tuning. Compared to existing supervised classification methods, the extraction of cell type knowledge in ItClust is tailored more towards the target data, which makes it more effective than other methods.

In summary, ItClust has shown to be a powerful tool for scRNA-seq clustering and cell type classification. It can accurately extract information from source data and apply it to help cluster cells in target data. It is robust to strong batch effect between source and target data, and is able to separate unseen cell types in the target. Furthermore, it provides confidence scores that facilitates cell type assignment. With the increasing popularity of scRNA-seq in biomedical research, we expect ItClust will make better utilization of the vast amount of existing well-annotated scRNA-seq datasets, and enable researchers to accurately cluster and annotate cells in their studies.

## Acknowledgements

This work was supported by the following grants: NIH R01GM108600, R01GM125301, R01HL113147, and R01EY030192 (to M.L), R01DK076077 (to. K.S.), and NSFC 31970649 (to G.H.).

## Author contributions

This study was conceived of and led by M.L.. J.H., X.L., G.H., and M.L. designed the model and algorithm. J. H. implemented the ItClust software and led the data analysis with input from M.L., X.L., G.H., Y.L., and K. S.. J.H. and M.L. wrote the paper with feedback from X.L., G.H., Y.L., and K.S..

## Competing financial interests

The authors declare no competing interests.

## Methods

### Information extraction from source data using the source network

ItClust requires two input datasets, a source dataset that includes cells with well-annotated cell type labels, and a target dataset that includes cells that need to be clustered and annotated. ItClust starts from selecting top *h* highly variable genes from the target data. Then, it finds the *p* overlapping genes in the source data (*p* ≤ *h*) and extracts the corresponding gene expression matrix for all *n* cells in the source data. Let *X* ∈ *R*^*n*×*p*^ be the extracted single-cell gene expression matrix used to train the source network in which rows correspond to cells and columns correspond to genes. Since scRNA-seq data are high-dimensional, instead of training directly in the original data space, we perform dimension reduction by transforming the data using a nonlinear mapping function *f_θ_*: *X* → *Z*, where *θ* represents embedding parameters, and *Z* ∈ *R*^*n*×*d*^ is the latent feature space, with *d* ≪ *p*. To parameterize *f_θ_*, we use a stacked autoencoder and initialize it layer by layer with each layer being a denoising autoencoder trained to reconstruct the previous layer’s output after random corruption [26]. We use ReLU as the activation function except for the bottleneck layer and last decoder layer, in which tanh is used as the activation function. After training each layer by minimizing the least-square loss, we concatenate all encoder layers followed by all decoder layers in reverse layer-wise training order to form a multilayer autoencoder, with a bottleneck layer in the middle, fine-tuning it to minimize the reconstruction loss. Then, we discard all decoder layers and use the encoder layers as our initial mapping between raw data and the dimension-reduced feature space.

Next, we add a clustering layer to the encoder network. Since the source dataset is labeled and the number of true cell types *k* is known, we use *k* to set the number of clusters. For each cluster *j*(*j* = 1,…, *k*), the cluster centroid *μ_j_* is determined by the mean features in the embedding layer based on cells in each cell type. This step assigns each cell type with an initial cluster and ensures that only cell type-specific gene expression signatures are captured during the later network optimization process.

### Transferring cell type information in the source data to the target network

Let *X*′ ∈ *R*^*m*×*p*^ be the gene count matrix for the target dataset with *m* cells, where *p* is a subset of the highly variable genes in the target data that are also present in the source data. We build a new network using the same structure as the source network. Rather than randomly initializing the target network, we transfer weights learned from the source network to the target network as initial values, except for the final clustering layer. This step ensures that the new network can map the target data to the same feature space *R^d^* as done in the source network, that is, *X*′ → *Z*′, where *Z*′ ∈ *R*^*m*×*d*^. In the next step, we apply an iterative clustering algorithm on the feature space *Z*′ [27], Since cell type labels for the target dataset are unknown, to get the number of clusters and cluster centroids, for each cell in the target dataset, we predict its cluster using the source network, and use the predicted number of clusters and mean features in each predicted cell cluster as the centroid to initialize the clustering layer.

### Fine-tuning of the target network

In addition to incorporating information from the source data, it is also necessary to incorporate unique information that is present in the target data only. To do so, we iteratively fine-tune the target network to make it more adaptive to the target data. During fine-tuning, for cell types that are present both in the source and target data, their cluster centroids will be mainly driven by the source data, and they will shift slightly to adapt to the target data over iterations. For cell types that are only present in the source data, none (or only a few) of the cells in the target data will be assigned to their centroids and we call these centroids as “free centroids”. For cell types that are unique to the target data, although there are no initial centroids allocated to them, those “free centroids” may move and are used to cluster those unseen cell types in the target data over iterations. Due to this reason, cell types that are unique in the source data can provide ItClust more flexibility and help cluster cell types that are unique in the target data. Thus, ItClust prefers a source dataset with comprehensive cell types, and such dataset can be obtained by combining multiple well-labeled datasets together.

### Parameter optimization for the source and target networks

The source and target networks are optimized using the deep embedding clustering algorithm [27]. First, we define the metric of distance from a cell to a cluster centroid. Following van der Maaten & Hinton [28], we use the Student’s *t*-distribution as a kernel to measure the distance between the embedded point *z_i_* = *f_θ_*(*x_i_*) ∈ *Z* for cell *i* and centroid *μ_j_* for cluster *j*:

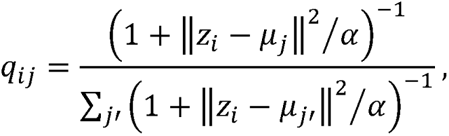

where *α* is the degree of freedom of the Student’s t-distribution. The distance *q_ij_* can also be interpreted as the probability of assigning cell *i* to cluster *j*.

Next, we iteratively refine the clusters by defining an auxiliary target distribution *P* based on *q_ij_*. The choice of *P* is important for ItClust’s performance. We define the auxiliary target distribution as:

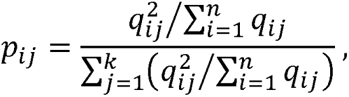

which upweighs cells assigned with high confidence, and normalizes the contribution of each centroid to the overall loss function to prevent large clusters from distorting the hidden feature space. Now that we have the soft assignment *q_ij_* and the auxiliary distribution *p_ij_*, we can define the objective function as a Ku I Iback-Leibler (KL) divergence loss:

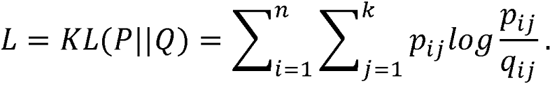

The network parameters and cluster centroids are simultaneously optimized by minimizing *L* using Stochastic Gradient Descent with momentum. The gradient of *L* with respect to *z_i_* and *μ_j_* are derived as:

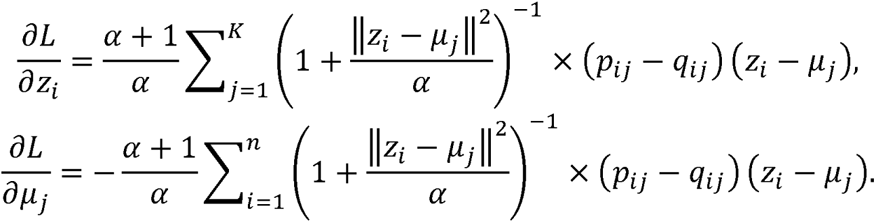

The gradients *∂L*/*∂z_i_* are used in standard backpropagation to calculate the network’s parameter gradients *∂L*/*∂θ* and *θL*/*∂μ_j_*, which are used to update the cluster centroids after finishing each batch with size *b* (for example, *b* = 256)by

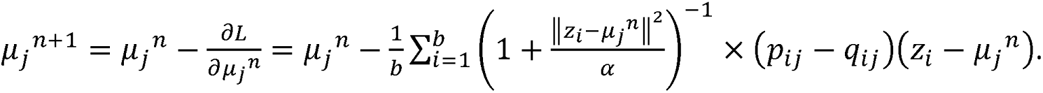

### Architecture of the neural network in ItClust

The number of layers and nodes for each layer depends on the sample size in the source data. Larger source data typically contain more information on cell types, and often require a larger network to store such information. We suggest different numbers of hidden layers and different numbers of nodes in the encoder. **Supplementary Table 2** gives the default numbers of hidden layers and nodes in ItClust.

### Cell type assignment

In addition to clustering, we also aim to leverage cell type information in the source data to help cell type assignment in the target data. Unlike most supervised cell type classification methods that predict cell type for each individual cell, ItClust provides a confidence score for each cluster to assist cell type annotation for all cells in that cluster. Since the source dataset is well-labeled, before fine-tuning, each cluster centroid in our model represents a known cell type present in the source data. For example, assume cluster *i* is used to cluster cell type *i* in the source data before fine-tuning. We first use the prefine-tuned model to assign clusters for cells in the target data. Let 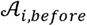 represent the set of cells in the target data that are assigned to cluster *i*. Since the pre-fine-tuned model was solely trained on the source data, cells in 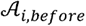 have gene expression patterns that resemble cell type *i* in the source data. During iterative fine-tuning, as the centroid for cluster *i* keeps updating its location, some cells in the target data may be added to and other cells may be removed from cluster *i*. After fine-tuning is finished, we get another set of cells 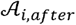, which consists all target cells that are assigned to cluster *i*.If the centroid for cluster *i* is still used to cluster cell type *i* in the target data, a big proportion of cells in set 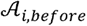 should also be present in 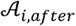. Based on this idea, a confidence score for each cluster can be calculated as the proportion of original cells in the final cluster:

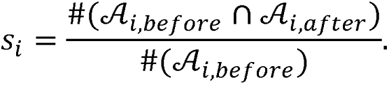

This confidence score measures how confident that cells in a given cluster in the target data are similar to a cell type that is present in the source data.

### Data preprocessing and quality control

ItClust can handle data in different formats including UMI count, FPKM, or TPM etc. All data follow the same preprocessing procedure. First, for both the source and target data, a cell is filtered out if the number of genes with non-zero expression is less than 100. We further remove MT and ERCC genes and genes that are expressed in less than 10 cells. The gene expression values are then normalized. In the first step, cell level normalization is performed in which each gene’s expression in each cell is divided by the total gene expression level in the cell, multiplied by 10,000, and then transformed to a natural log scale. In the second step, gene level normalization is performed in which the cell level normalized values for each gene are standardized by subtracting the mean across all cells and divided by the standard deviation across all cells for the given gene.

Highly variable genes are selected using the filter_genes_dispersion function from the Scanpy package [29] (https://github.com/theislab/scanpy). Selection of top highly variable genes is based on the target dataset only. By doing this, we ensure that all selected genes are informative to distinguish cells in the target data. Next, we find the overlap between the highly variable genes in the target data and those that are present in the source data, and we then extract the expression patterns of these overlapped genes from the source data.

### Downsampling experiments on UMI counts

We performed downsampling experiments only for datasets with UMI counts. To generate a downsampled dataset from the original scRNA-seq data, we selected high-quality cells and genes with high expression from the original dataset. The observed expression level of gene *g* for cell *c* is treated as the true expression *λ_gc_*. We generate downsampled datasets by drawing from a Poisson distribution with mean parameter *τ_c_λ_gc_*, where *τ_c_* is the cell-specific efficiency. This ensures the downsampled dataset and the original dataset are similar in mean expression and the percentage of zero entries. To mimic variation in efficiency across cells, we sampled *τ_c_* as follows, 75% efficiency with *τ_c_~Gamma*(10,0.075), 50% efficiency with *τ_c_~Gamma*(10,0.0.05), 25% efficiency with *τ_c_~Gamma*(10,0.025), and 10% efficiency with *τ_c_~Gamma*(10,0.01).

### Datasets

We analyzed multiple scRNA-seq datasets. Publicly available data were acquired from the access numbers provided by the original publications. Details of the datasets analyzed in this paper were described in **Supplementary Table 1.**

### Software availability

An open-source implementation of the ItClust algorithm can be downloaded from https://github.com/jianhuupenn/ItClust

### Life sciences reporting summary

Further information on experimental design is available in the Life Sciences Reporting Summary.

