## Supplementary Materials for "Iterative transfer learning with neural network for clustering and cell type classification in single-cell RNA-seq analysis"

**Supplementary Table 1. Datasets analyzed in this paper.**

| **Dataset** | **Organism** | **Tissue** | **Number of cells** | **Experimental protocol** | **Cell types (number of cells)** |
| --- | --- | --- | --- | --- | --- |
| Baron *et al*. | Mouse | Pancreas | 1,886 | InDrop | beta (894); ductal (275); delta (218); alpha (191); endothelial (139); quiescent stellate (47); gamma (41); macrophage (36); activated stellate (14); B cell (10); immune other (8); T cell (7); schwann (6). |
| Baron *et al*. | Human | Pancreas | 8,569 | InDrop | beta (2,525); alpha (2,326);  ductal (1,077); acinar (958);  delta (601); activated stellate (284); gamma (255); endothelial (252); quiescent stellate (173); macrophage (55); mast (25); epsilon (18); schwann (13); T cell (7). |
| Xin *et al*. | Human | Pancreas | 1,492 | SMARTer | alpha (886); beta (472); gamma (85); delta (49). |
| Grün *et al*. | Human | Pancreas | 1,004 | CelSeq | ductal (327); acinar (228);  alpha (191); beta (161);  delta (50); activated stellate (19); gamma (18); endothelial (5); schwann (1); quiescent stellate (1); mast (1); macrophage (1); epsilon (1). |
| Muraro *et al*. | Human | Pancreas | 2,285 | CelSeq2 | alpha (843); beta (445); acinar (274); ductal (258); delta (203); gamma (110); activated stellate (90); endothelial (21); macrophage (15); quiescent stellate (12); mast (6); schwann (4); epsilon (4). |
| Lawlor *et al*. | Human | Pancreas | 638 | Fluidigm C1 | beta (258); alpha (239); ductal (36); delta (25); acinar (21); gamma (18); activated stellate (16); endothelial (14); schwann (5); mast (3); quiescent stellate (1); macrophage (1); epsilon (1). |
| Segerstolpe *et al.* | Human | Pancreas | 2,394 | Smart-Seq2 | alpha (1,008); ductal (444); beta (308); gamma (213); acinar (188); delta (127); activated stellate (55); endothelial (21); epsilon (8);  Mast (7); macrophage (7); quiescent stellate (6); schwann (2). |
| Park *et al*. | Mouse | Kidney | 43,745 | 10X | PT (26,482); DTC (8,544); CD-IC (1,729); LOH (1,581); T lymph (1,308); Endo (1,001); CD-PC (870); Novel (643); Fib (549); NK (313); B lymph (235); Macro (228); CD-Trans (110); Podo (78); Neutro (74). |
| Our own data | Human | Kidney | 15,693 | 10X | PT (3,695); Macrophage (3,566); T Cells (2,721); Endo (1,837); NK_Cells (1,430); CD_IC (1,297); Loop of Henle (551); Distal Tubules (383); B Cells (213). |
| Peng *et al*. | Macaques 1,2, and 3 | Retina | 25,597 | Drop-seq | IMB (5,066); FMB (4,457);  RB (2,814); DB5* (2,775); DB3b (2,538); DB4 (2,340); DB2 (1953); BB/GB* (1488); DB1 (935); DB3a (605); DB6 (547); OFFx (79). |
| Peng *et al*. | Macaque 4 | Retina | 4,705 | Drop-seq | RB (1262); IMB (1085); DB5* (692); DB4 (645); BB/GB* (327); DB2 (291); DB6 (111); DB3b (102); OFFx (68); DB1 (61); FMB (43); DB3a (18). |

**Supplementary Table 2. Default hyperparameters of the autoencoder.**

| **No. of cells in source data** | **No. of hidden layers** | **No. of nodes in the 1st hidden layer** | **No. of nodes in the 2nd hidden layer** |
| --- | --- | --- | --- |
| >10,000 | 2 | 128 (or larger) | 32 |
| (5,000,10,000] | 2 | 64 | 32 |
| (2,000,5,000] | 2 | 64 | 32 |
| (500,2,000] | 1 | 64 | 0 |
| <500 | 1 | 16 | 0 |

**Supplementary Table 3. Programs compared in this paper.**

| **Method** | **Software** | **Version** | **Url** | **Reference** |
| --- | --- | --- | --- | --- |
| DESC | desc | 1.0.0.5 | https://github.com/eleozzr/desc | [1] |
| Seurat3.0 | Seurat | 3.0.0 | https://github.com/satijalab/seurat | [2] |
| Louvain | scanpy | 1.3.1 | https://icb-scanpy.readthedocs-hosted.com/en/stable/api/scanpy.tl.louvain.html | [3] |
| Moana | Moana | 0.1.1 | https://github.com/yanailab/moana | [4] |
| scmap | scmap | 1.1.6 | https://github.com/hemberg-lab/scmap | [5] |
| SAVER-X | SAVER-X | 1.1.2 | https://github.com/jingshuw/SAVERX | [6] |

**Supplementary Note: Analysis of the human kidney data**

**Human kidney data (target dataset)**

Dataset 1: This dataset includes 35,908 cells and 32,738 genes from 4 normal human kidneys, generated by us using 10X.

Dataset 2: This dataset was generated by Young *et al* [7]. Single-cell transcriptomes from human kidneys reveal the cellular identity of renal tumors. Science 361(6402):594-599.

The dataset was download from Data S1 (<http://science.sciencemag.org/highwire/filestream/713964/field_highwire_adjunct_files/4/aat1699_DataS1.gz.zip>) in the Supplementary Materials of Young *et al*. (2018). In this analysis, we focused on normal kidney cells (total 10,621 cells), which are from VHL (2,706 cells), RCC1 (3,747 cells) and RCC2 (4,168 cells).

We combined Dataset 1 and Dataset 2 in the analysis. The resulting data include 46,529 cells, and 31,232 shared genes between these two datasets. The combined data can be downloaded from <https://www.dropbox.com/s/l9zq2sge93n4ifj/human_kidney_desc_use.tar.gz?dl=0>.

Cell filtering criteria: 1) eliminated cells with percentage of mitochondrial UMI counts >20%; 2) eliminated cells with gene counts <200; 3) eliminated cells with total UMI counts <1,000.

Gene filtering criteria: 1) eliminated genes if the number of cells expressing this gene is <10.

Data processing: 1) gene expression levels for each cell was normalized using the “scanpy.api.normalize_per_cell” function in scanpy with counts_per_cell_after =10,000; 2) top 1,000 highly variable genes were selected using the “scanpy.api.pp.filter_genes_dispersion” function in scanpy; 3) normalized gene expression for the selected top 1,000 highly variable genes was then transformed using log(1+x) transformation with natural logarithm; 4) the expression is further standardized to a z-score, and the standardized gene expression values were used as input for DESC [1]. After the above filtering and data processing, there were 15,693 cells×1,000 highly variable genes remained in downstream analysis.

Clustering and identification of cell types: All 15,693 cells were clustered by DESC [1]. We used two hidden layers with 128 nodes in the first hidden layer, and 32 nodes in the second hidden layer. And resolution is set to 0.4. The clusters obtained from DESC were labeled by marker genes provided by Young *et al*. [7].

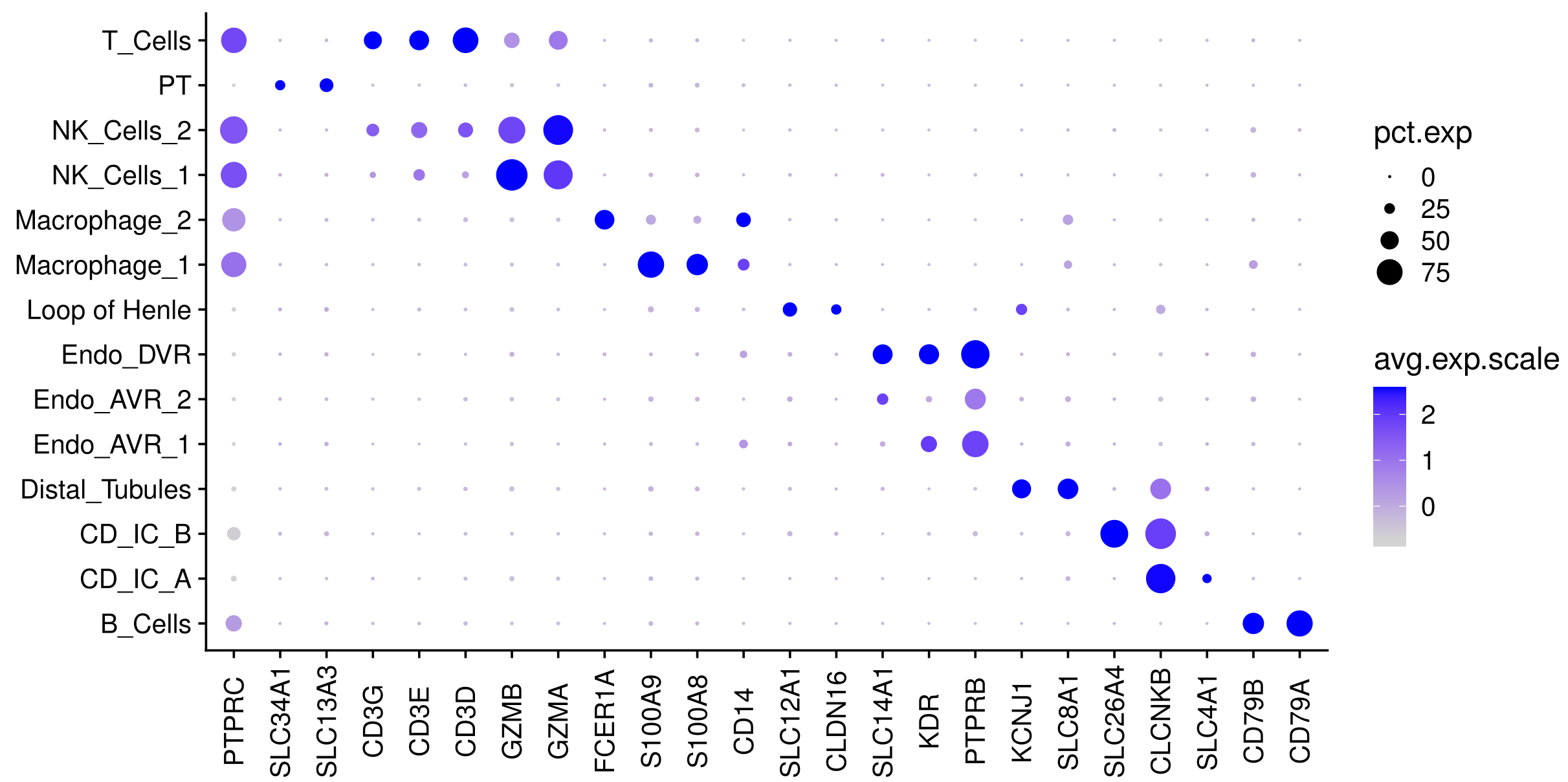

**Supplementary Fig 1.** Dot plots of known marker genes used for cell type identification for the human kidney data (data generated ourselves together with data from Young et al. [22]). The marker genes used to label the cell types are: *SLC13A3 and SLC34A1 for PT (Proximal Tubule); CLDN16 and SLC12A for Loop of Henle; PTPRB and KDR for Endo_AVR_1 (Endothelial Ascending Vasa Recta); PTPRB and SLC14A1 for Endo_AVR_2; PTPRB, KDR, and SLC14A1 for Endo_DVR (Endothelial Descending Vasa Recta); KCNJ1 and SLC8A1 for Distal Tubules; SLC4A1 and CLCNKB for CD_IC_A; SLC26A4 and CLCNKB for CD_IC_B; GZMA and GZMB for NK_cells; CD3D, CD3E, and CD3G for T_cells; CD14, S100A8, and S100A9 for Macrophage_1; CD14 and FCER1A for Macrophage_2; CD79A and CD79B for B_cells.*.

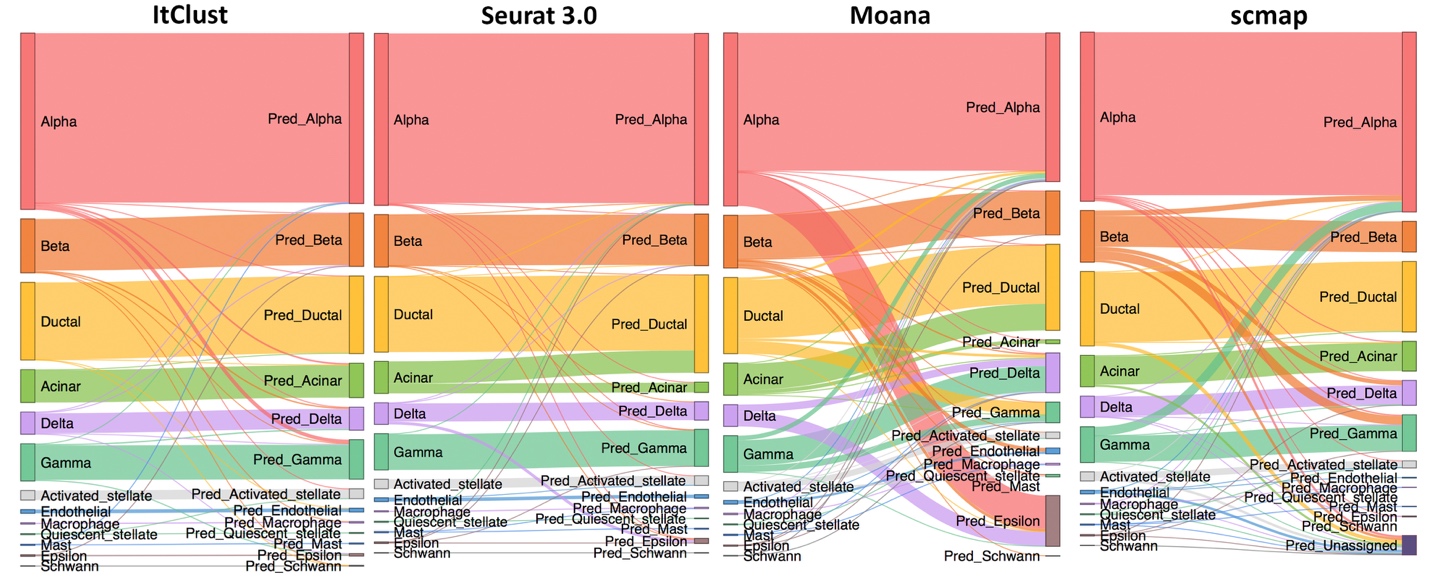

**Supplementary Fig 2.** The Sankey plots of ItClust, Seurat 3.0, Moana and scmap cell type classification results for the Segerstolpe *et al*. dataset using the combined source data.
