## Supplementary figures and images for "Iterative transfer learning with neural network for clustering and cell type classification in single-cell RNA-seq analysis"

### Supplemental Figure 1,

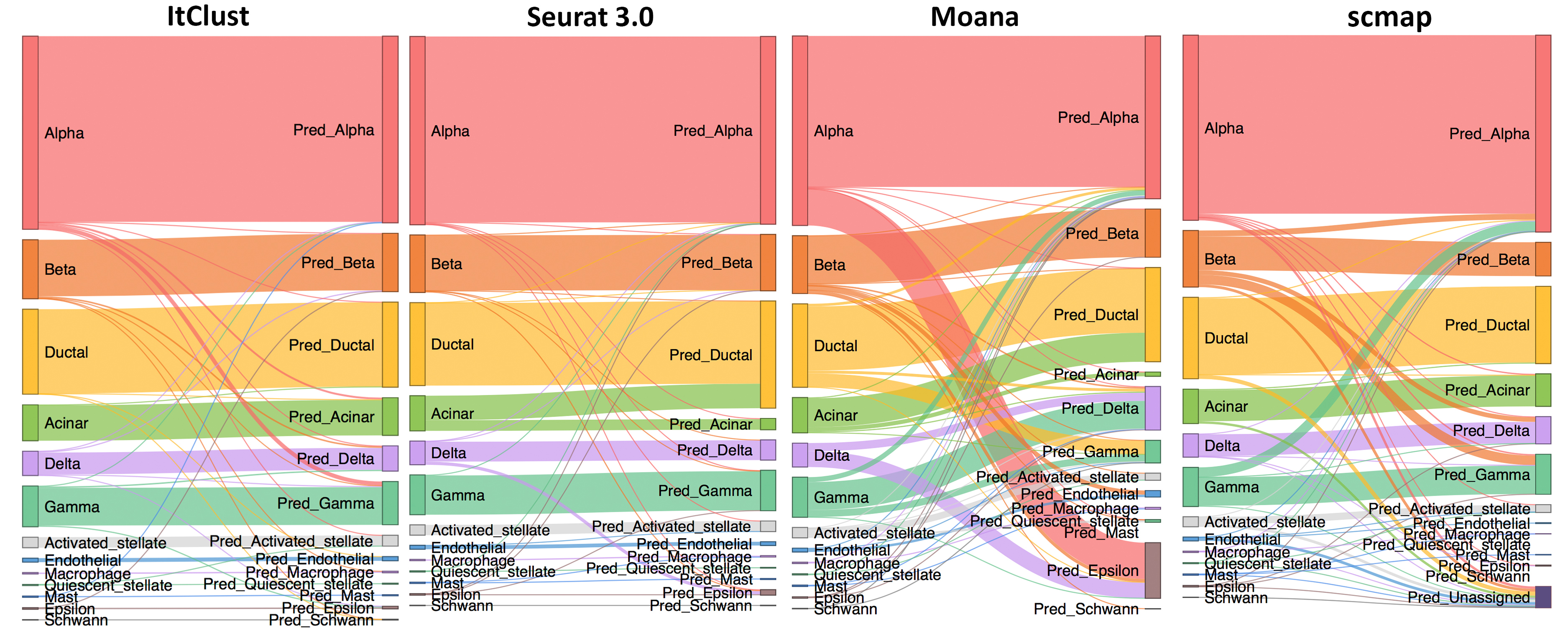
